# The microbiota elicits compensatory adaptation in a seasonally-adapting animal host

**DOI:** 10.1101/2025.10.14.682128

**Authors:** Dean B. Peterson, Rebecca Kreutz, Peyton Jackson, Hyrum O. Pech-Gonzalez, J. Kelly Moon, Jacky Chan, Susan Wilcox, Jack Beltz, Paul Schmidt, John M. Chaston

**Affiliations:** Department of Plant and Wildlife Sciences, Brigham Young University, Provo, Utah, USA, 84602; Department of Biology, University of Pennsylvania, Philadelphia, PA 19104

**Keywords:** *Acetobacter*, *Weissella*, local adaptation, life history, rapid adaptation

## Abstract

Seasonal adaptation in *Drosophila melanogaster* is a model for understanding the evolutionary responses of organisms to cyclical environmental changes, including roles played by associated microorganisms (‘microbiota’). Here we examined how the microbiota influences *D. melanogaster* seasonal adaptation by rearing flies in outdoor mesocosms, fed diets inoculated with different bacterial strains that have distinct influences on the flies’ life history. The bacterial treatments influenced fly population dynamics and microbiota composition over a summer-to-fall season. The developmental phenotype of the treated flies initially differed but converged over time in flies reared with a complete microbial community. Conversely, rearing the flies free of their colonizing microorganisms revealed that the bacterial treatments led to evolution of distinct developmental phenotypes. The development time of flies from the different treatments consistently adapted to compensate for the direct influence of the bacteria on host development; e.g., flies evolved faster development times if they were inoculated with microbes that slowed development. This compensatory trend was apparent in flies reared in a second location and season, and is consistent with a previous report of wild-sampled flies whose development phenotype segregated with their microbiota composition. Together, these results reveal that microbiota-dependent selection consistently elicits compensatory adaptation in seasonally-evolving flies, which we conclude is a mechanism whereby horizontally-acquired, low-fidelity microbial partners can shape the evolution of their animal hosts.

**Importance:** Understanding how organisms adapt to seasonal change can model evolutionary responses in a broader changing world. *Drosophila melanogaster* is a powerful model for studying these dynamics, especially when considering the influence of transient yet impactful colonizing microorganisms. This study reveals that microbial partners can drive consistent, compensatory adaptation in host development across seasons and locations. By demonstrating that flies evolve faster development times in response to microbes that slow a model host’s period of growth and development, this work highlights one way that the microbiota can influence adaptation in their animal hosts. These findings also provide evidence that low-fidelity, horizontally-acquired microbes can exert selective pressures strong enough to shape host life history traits. These insights underscore the microbiota’s role as an ecological and evolutionary force.

## Introduction

Understanding how organisms adapt to their environments is a central question in evolutionary biology. While much traditional focus has been on genetic changes driven by abiotic factors including temperature, humidity, and resource availability, recent advances have highlighted the profound influence of biotic interactions, as with microbial symbionts, and their likelihood to shape evolutionary trajectories of their hosts (1-4). This expectation underscores the increasing importance of the microbiota not merely as passengers but as dynamic agents capable of influencing host fitness and adaptation, in order to more completely understand the spectrum of evolutionary pressures that act on animals (5-11).

Animal adaptation occurs through a variety of mechanisms, including mutation, recombination, and selection. Traditionally, these processes have been studied in the context of environmental stressors or ecological competition. However, the microbiota can also act as a selective force, influencing host traits and evolutionary outcomes. Microbial communities can modulate nutrient acquisition, responses to infection, stress tolerance, and developmental signaling pathways, thereby affecting host survival and reproductive success (12-16). In some cases, maladaptive host genotypes may persist if microbial partners compensate for genetic deficiencies. This can lead in the extreme to host-microbe interdependence, as in the nutritional dependencies of certain insects and their vertically-transmitted symbionts (17-19). Transformative insights into co-adaptive and arms-race interactions have and will come from such studies relative to obligate symbionts or pathogens, but the interactions between low-fidelity partners, including so-called commensals and their hosts, are not as well understood (1, 20-22). A clearer picture of how commensals influence host adaptation would not only reinforce how the microbiota can be a driver of host genetic selection, but predict the outcomes of the microbial influences.

Powerful experimental frameworks have been established for investigating adaptation in *D. melanogaster*, including through the use of outdoor mesocosms that expose flies to various seasonal pressures but prevent gene flow so that genomic variation in the experiment can be attributed to seasonal change (23). Outdoor mesocosms provide a unique opportunity to observe evolutionary processes as they unfold in real time, capturing the interplay between environmental pressures, microbial interactions, and host genetic variation. These systems bridge the gap between laboratory precision and ecological relevance, offering insights that are difficult to obtain in either setting alone. Experiments using these approaches have established that, among other things, the microbiota can be an agent of host genetic selection over relatively short timescales measured in weeks (24) or months (25).

In addition to being a model for adaptation to a changing environment, the microbiota of *Drosophila* is useful for exploring beneficial host-microbe associations in the laboratory and in wild settings (26, 27). The microbiota can influence a wide range of host life history traits including nutrient storage, development rate, fecundity, stress resistance, and lifespan (28-33). Dominant members of the microbial communities usually include Lactic Acid Bacteria (LAB), Acetic Acid Bacteria (AAB), and gamma-Proteobacteria (34-41). Different strains in the community can have markedly different influences on host phenotypes in monoassociation, and *D. melanogaster* species collected from different geographic locations genetically select for distinct microbial communities (28, 42). Additionally, the composition of the microbiota can shift in response to environmental conditions, creating a dynamic feedback loop between host genotype, microbial community structure, and ecological context (25, 43-45). These interactions suggest the capacity of the microbiota to play a key role in host adaptation, particularly under conditions of genetic or environmental stress. We recently observed that the phenotypes of in wild flies bearing distinct microbial communities had a distinct relationship to the inferred functions of their microbial communities; that is, that hosts bearing abundantly bacteria that are expected to accelerate host development tended to have slower evolved development times (44). Conversely, hosts with higher loads of bacteria that normally slow host development had evolved faster development times. We suggested that these patterns were consistent with compensatory influences of the microbiota on host adaptation. However, such patterns have not yet been demonstrated experimentally.

In this study, we examine whether the microbiota elicits compensatory evolution in *Drosophila melanogaster* reared in outdoor mesocosms. By comparing host genotypes across microbiota treatments and environmental conditions, we show that different microbial strains can enhance or buffer the developmental phenotypes of wild-adapting flies, and that the flies adapt in response to these fitness influences of their microorganisms. Our approach integrates ecological reality with evolutionary analysis, providing a framework to explore how microbial symbionts influence host adaptation in complex environments. Ultimately, this work contributes to a growing body of evidence that the microbiota is not only a modulator of host phenotype but also a potent force in shaping host evolutionary outcomes.

## RESULTS

*The microbiota shapes* D. melanogaster *population traits over a summer-to-fall season* We measured the microbial influence on *D. melanogaster* seasonal adaptation by continuously exposing flies in replicated outdoor mesocosms to diets inoculated with one of *Acetobacter orientalis* DmW_045 (*Ao*), *Acetobacter* sp. SLV-7 DmW_125 (*As*), or *Weissella paramesenteroides* DmW_115 (*Wp*). Census estimates over ∼ 120 days revealed that population size varied significantly with time (χ^2^ _10, 1302_ = 602.65, p < 10^−15^) and bacterial treatment (χ^2^ _2, 1302_ = 15.49, p < 0.0004). As has been reported for flies in the wild or in similar mesocosm setups at other locations absent of bacterial inoculations (24, 46, 47), population size peaked during the summer and declined with onset of winter (**FIG 1A**). The most striking pattern was that flies inoculated with *As* had better survival during a late-season temperature spike, and that their populations remained higher as cold temperatures onset. Together, these findings reveal that the flies in our conditions follow the same general trends as other outdoor mesocosm experiments with fruit flies, and that season-long inoculation with specific bacterial species altered the flies’ population sizes over time.

**Fig 1.**
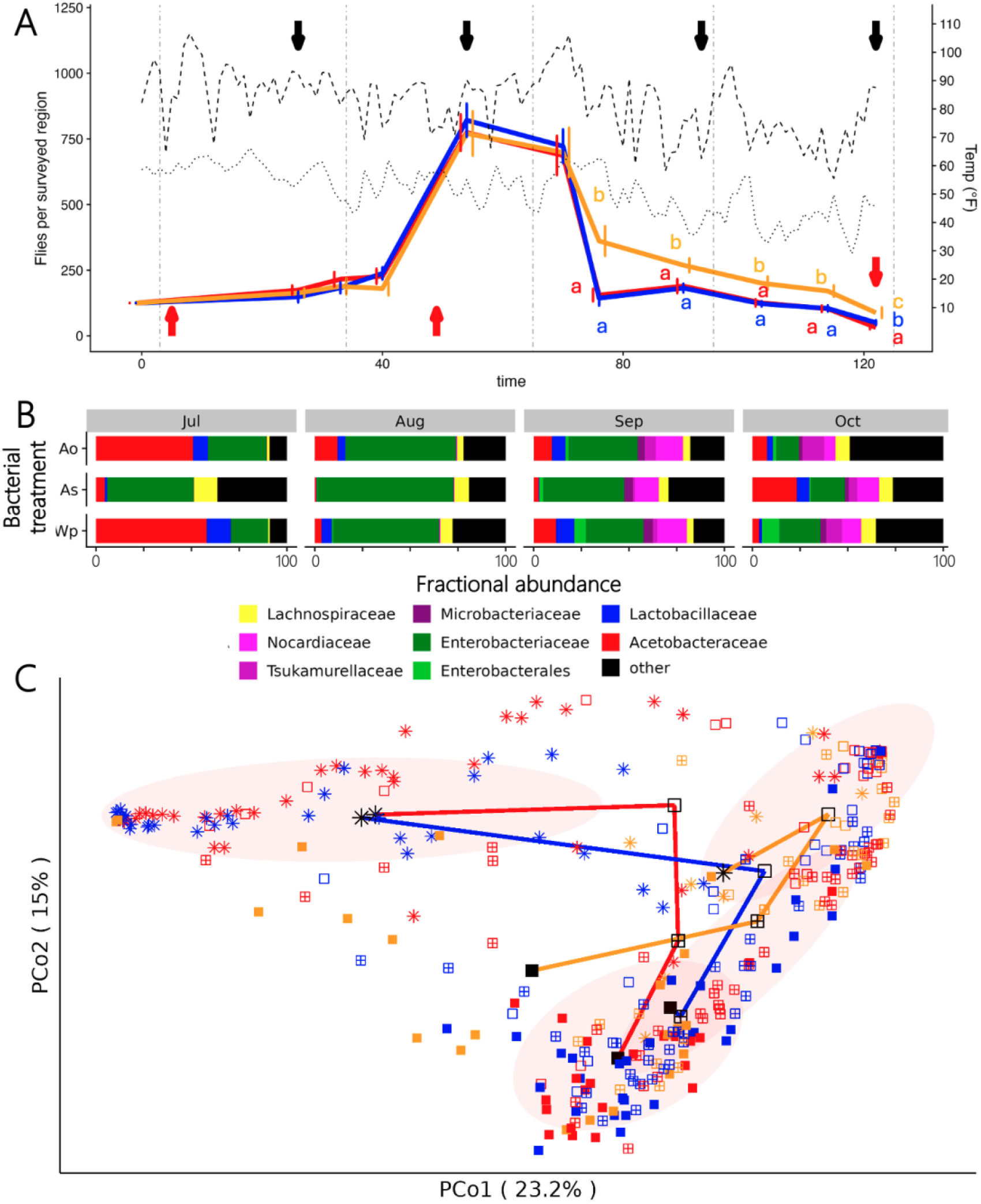
Seasonal adaptation observed through population. A) Fly population sizes (solid lines) and daily maximum (dashed line) and minimum (dotted line) temperatures were measured in flies reared in outdoor mesocosms. Compact letter displays are colored according to bacterial treatment and are present whenever there was a significant effect of the bacterial treatment on population size at a given time. Black arrows indicate when flies were sampled for microbiota analysis and red arrows when flies were sampled and raised in the lab for phenotypic analysis. Vertical lines delineate months. B) Bacterial genera are indicated by different colors, and the plot is separated into panels representing combinations of date (July, Aug, Sep, Oct) and treatment (*Ao, As, Wp*). C) Principal coordinates are separated by treatment (red, *Ao;* orange, *As;* blue, *Wp*) and by date (star, July; open square, Aug; hatched square, Sep; filled square, Oct).

To better understand the pressures that influenced the adapting fly populations, we characterized the flies’ microbiota. Consistent with most analyses of the fly microbiota in the wild, the most abundant bacterial families in the flies were from the gamma-proteobacteria, LAB, and AAB (**FIG 1B**). The flies’ microbiota composition varied with sampling date (F _3, 317_ = 30.26, R^2^ = 0.20, p < 0.001), microbial treatment (F _2, 317_ = 3.98, R^2^ = 0.02, p < 0.001), and the interaction of the two (F _6, 317_ = 3.55, R^2^ = 0.05, p < 0.001), (**TABLE S1**). Microbiota composition also varied with outdoor mesocosms the flies were collected from (F _12, 317_ = 2.75, R^2^ = 0.07, p = 0.001). Principle Coordinate Analysis (PCoA) showed that relationship between the microbiota composition of the different treatments mirrored the treatment patterns for fly population size. As with fly population size, the starting and ending microbiota composition of flies inoculated with *As* tended to be distinct from the *Ao-* and *Wp-* inoculated flies, as did the overall trajectory of microbiota change over time (**FIG 1C**). Many different microbial taxa contributed to seasonal variation in microbiota composition, including *Wolbachia* (abundance declined over time) and fourteen other bacterial families (**FIG S1**). The *Acetobacteraceae* and *Lactobacillales* were each most abundant in *Ao*- or *Wp*-inoculated flies when the experiment began, and in *As*-inoculated flies at the experiment’s conclusion. Only two family-level groups, reads assigned to the *Lactobacillaceae* and the Enterobacteriales, varied with the microbial treatment applied to the mesocosms (**FIG S1D**,**H**). Taken together, the correlated shifts in fly population sizes, overall microbiota composition, and the abundances of these major microbial groups over time together suggest a link between the microorganisms and fly phenotypes in a wild setting.

*The microbiota shapes* D. melanogaster *adaptation over a summer-to-fall season* We next sought to understand how the bacterial inoculation influenced the flies’ rapid adaptation by comparing life history and microbiome traits of the evolved flies. We measured traits in flies taken from the field and reared in the laboratory under bacteria-controlled conditions. The rationale for these experiments is that the phenotypes of axenic (‘bacteria-free’) flies reflect their genetic adaptations independent of any microbial influence, and the phenotypes of gnotobiotic flies show the flies’ traits absent confounders of varied microbial sampling. Fly development time and microbiota composition varied with time and bacterial treatment in gnotobiotic flies and, for development time, axenic flies (**FIG 2**). *As*-inoculated flies evolved to select for a lower fractional, but not absolute, LAB abundance (**FIG 2G-I**). Also, by the conclusion of the experiment there were different evolved differences in development time (**FIG 2A**) that were masked by the presence of the microbial community (**FIG 2B**). By October, the flies had evolved axenic differences in development time that were the reciprocal of the direct effect of the inoculated microorganisms on a trait (**FIG 2A,C**); in other words, *Ao-*inoculated flies had evolved the slowest development times of the three treatments even though *Ao* confers faster development times on flies than either *As* or *Wp*. These host-programmed differences disappeared in flies reared with a multi-species microbial community (**FIG 2B**), showing that the adaptive patterns in the axenic flies interacted with the microbial composition of the flies to converge on a development time phenotype that likely represents the field-selected value of the trait. Unlike development time, fly starvation resistance varied with time but not with bacterial treatment (**FIG 2 B,E**). And, fly fecundity, which was not influenced by the strains we examined, did not vary with time or with bacterial treatment (**FIG S2**). Together, these experiments suggest that the flies’ development rates and genetic control of the microbiota may be particularly susceptible to microbiota-dependent selection on the host, and that evolved host genetic differences can interact with the microbiota to determine host phenotypes.

**Fig 2.**
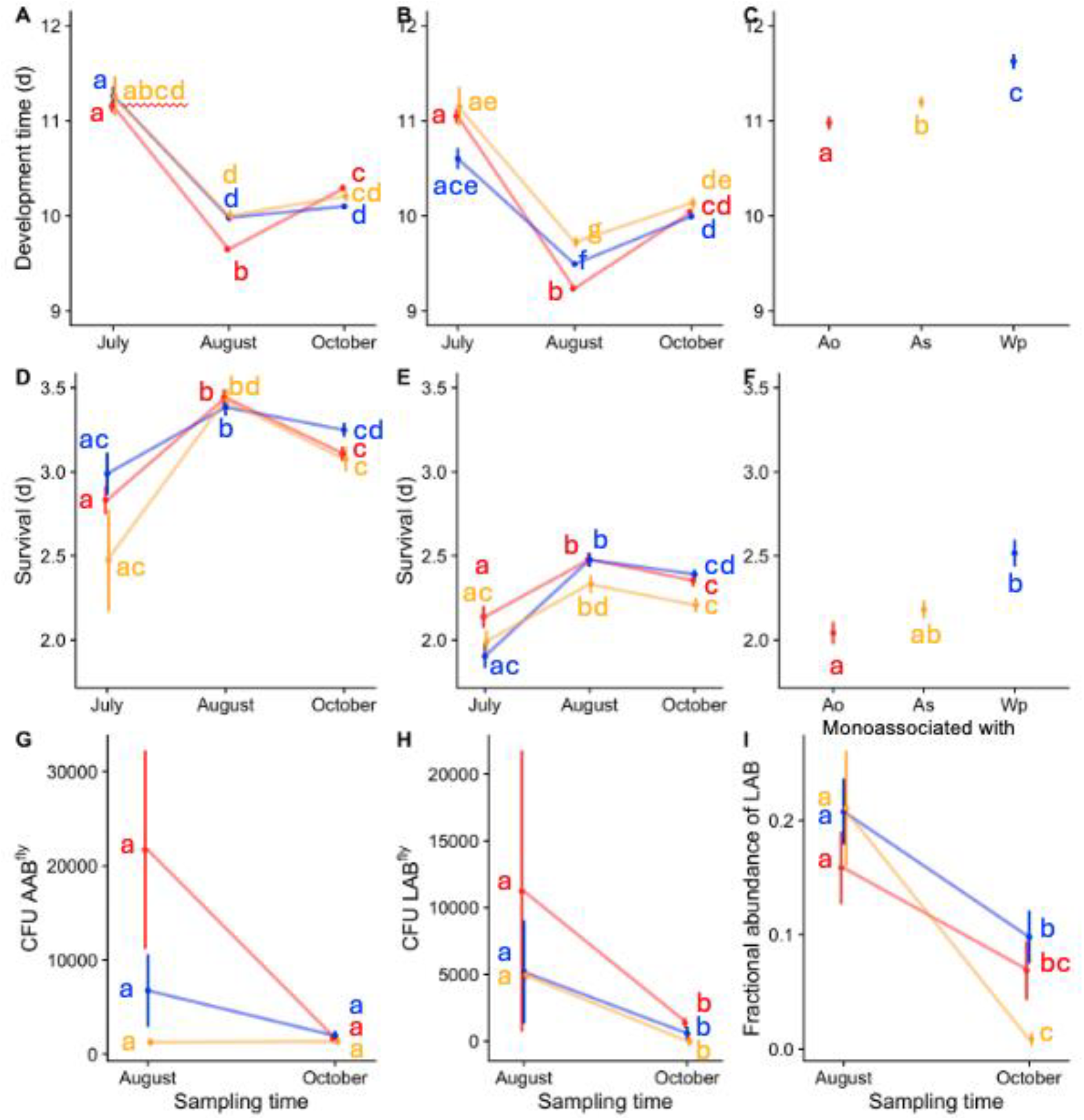
Life history traits of these seasonally adapting fly populations. Flies inoculated with *Ao* (red), *As* (orange), or *Wp* (blue) in the field were reared in the laboratory under axenic (A,D), 6-species gnotobiotic (D, E, G-I), or monoassociated (C,F) conditions. Flies were sampled in Aug, Oct, and sometimes July, reared under the specific conditions in the laboratory, and their A-C) development time, D-F) starvation resistance, and G-I) microbiota composition was measured. Flies in C and F were a pool of flies from all mesocosms, collected from the starting July population. Compact letter displays identify significant differences as determined by a Cox proportional hazards model (A-F) or a Kruskal-Wallis test (G-I).

*There are replicated evidences of compensatory seasonal evolution in* D. melanogaster Finally, we compared our results with published (44) and unpublished findings from other studies of microbiota-dependent evolution of fruit flies in a wild setting. Development time of the flies in this study evolved to compensate for the direct influence of the microorganisms. The same microbiota-compensatory pattern of host adaptation was evident in the development times and starvation resistances of flies that had been evolved in outdoor mesocosms at Pennsylvania, USA when inoculated with either *A. tropicalis* or *L. brevis* (data are newly reported from flies studied in (24), **FIG 3**). When reared axenic in the laboratory, the flies evolved with *A. tropicalis* took longer to develop to reproductive maturity and survived longer under starvation conditions than flies evolved with *L. brevis* (**FIG 3A,C**). However, the direct effect of *A. tropicalis* on that fly population, relative to *L. brevis*, was to shorten the time periods of these traits (**FIG 3B,D**). We also previously reported the same compensatory pattern in wild flies collected from a commercial orchard at Maine, USA from apples (which lead the flies to have AAB-dominated microbial communities) or compost (which increases the relative abundance of LAB in wild flies): flies from sources inoculated with or leading to higher levels of AAB had longer axenic development times than flies inoculated with LAB or reared in LAB-promoting conditions (44). Taken together, a consistent outcome of these experiments in different conditions, environments, and timescales is that the phenotypic patterns of the host evidence compensatory evolution relative to the effects of the environmental microorganisms.

**Fig 3.**
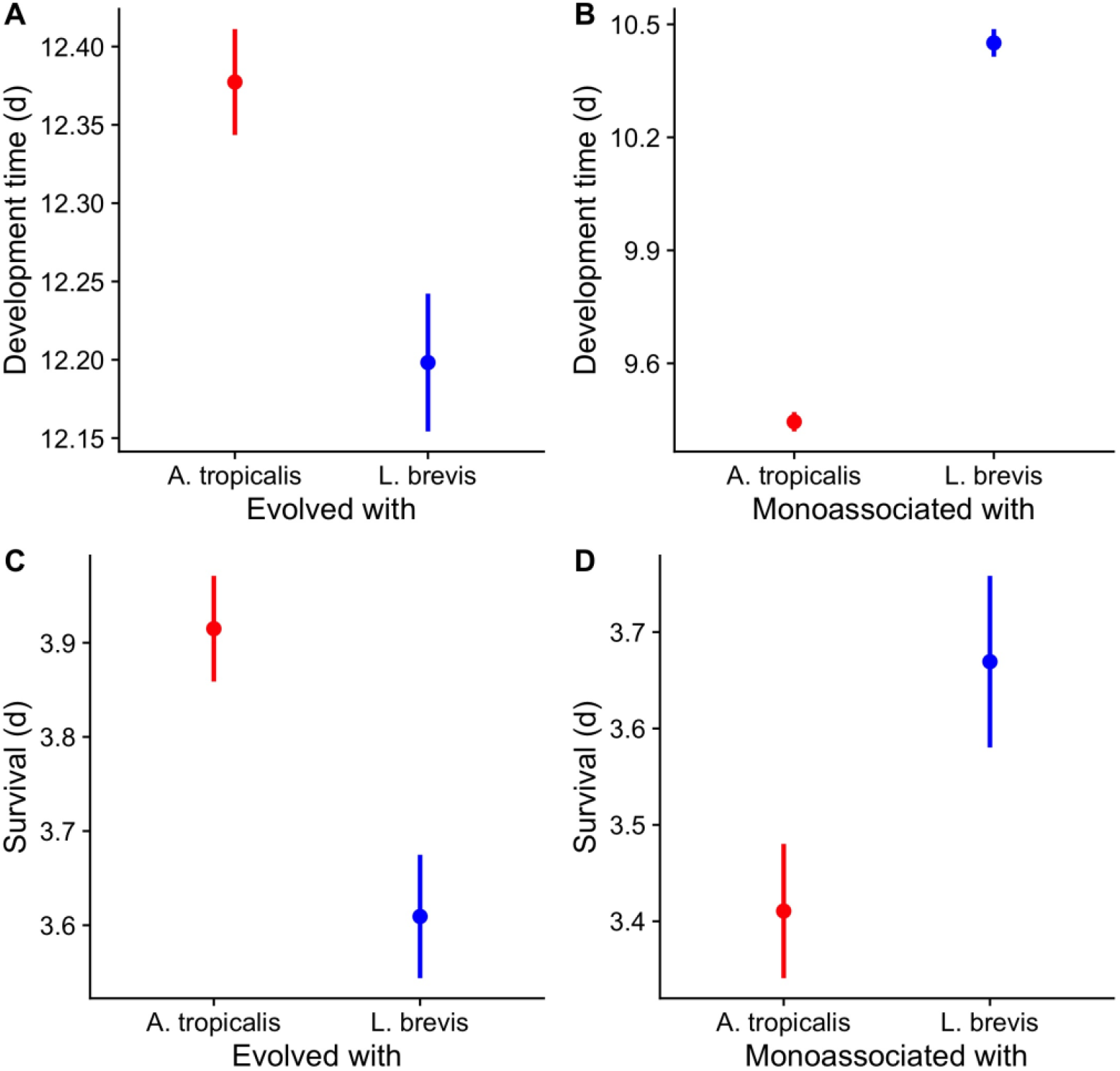
Life history traits of a second set of seasonally adapting fly populations. Flies inoculated with *A. tropicalis* (red), or *L. brevis* (blue) in field mesocosms in Media, PA as reported previously (24) were reared in the laboratory under axenic (A,C) or monoassociated (B,D) conditions after 6.5 weeks of outdoor selection. In every panel, there were significant differences between flies evolved or monoassociated with *A. tropicalis* and *L. brevis* as determined by a Cox proportional hazards model.

## DISCUSSION

We provide evidence here that microbial exposure can shape the seasonal adaptation of *Drosophila melanogaster* in naturalistic settings. By continuously inoculating outdoor mesocosms with distinct bacterial strains, we observed that microbial identity significantly influenced fly population dynamics, microbiota composition, and host-controlled host development times. Along with host-programmed starvation resistance these traits also all varied with season. These findings build on previous mesocosm studies (24) by demonstrating that microbial associations are not only responsive to environmental conditions but can actively mediate host adaptation across seasonal transitions. Together with previous analyses, we also find a consistent signal of compensatory *D. melanogaster* evolution in response to these varied microbial influences.

Our interpretation that compensatory evolution is a feature of host responses to the microbiota comes with several caveats. Among these, the most significant is the timing when compensatory evolution is evident. In our main experiments, compensatory evolution was apparent only in the axenic development times of the flies exposed to different microorganisms at the conclusion of the experiment, simultaneous with convergence of the development time phenotype across the flies (comparing their gnotobiotic, not their axenic, phenotypes in the laboratory). Conversely, the flies at the summer sampling displayed cooperative adaptation, where inoculation with *Ao*, which confers the shortest development times on the flies, led to the shortest host genotype-determined development times of the three treatment conditions. This observation provides two additional insights. First, while compensatory adaptation was more commonly observed (in each of three different studies identified here), microbiota-cooperative adaptation can also occur. Second, microbiota-compensating host adaptation is season independent because the Philadelphia populations that evolved only during the summer showed the same patterns as end-of-season populations in this study. We suggest that differences in the location, genotypes, and diets of the fly populations in the two separate experiments, and one or more of these factors could have affected the different timing of selection outcomes.

Separately, our interpretation of compensatory adaptation in wild flies collected from two dietary substrates in Maine comes with the caveat that we assume LAB strains slow and enhance selection for faster development time of the flies. This conclusion is consistent with the general effects of LAB strains used in our two selection studies, but we cannot rule out that the strains in the flies at the time did not meet this expectation. Whereas the evolution of development rate had similar outcomes in two experimental evolution studies, the SR outcomes of flies reared in Pennsylvania were not the same as in our study (SR was not measured in flies collected at Maine, USA). As asserted above, the discrepancy between these outcomes could be due to any number of varying factors, including the starting genotype of the flies, the wider difference in phenotypes conferred by *A. tropicalis* and *L. brevis* relative to *Ao, As*, and *Wp*, or differences in the environmental or experimental (e.g. dietary composition) features of the two experiments. Regardless of these varied influences, which provide substantial space to expand our understanding of whether specific environments, conditions, or other selective pressures potentiate this pattern, compensatory host selection was the most common outcome observed or inferred across these studies.

Inoculating the flies with the microbes substantially influenced microbiota composition and fly phenotypes, yet unexpectedly the inoculated strains were rare and low abundance in adult flies. Indeed, our initial rationale for selecting these strains was that *Wp* and *Ao* both colonize the flies at high levels even under conditions where flies are frequently transferred to new diets; and that *As* is a poor long-term colonizer (48). Our previous experimental evolution work had used two strains with only modest long-term colonization phenotypes, and we had anticipated that using strong long-term colonizers would lead to high, long-term fly colonization. Instead, the *Wp* and *Ao* strains were both very rare and very low in abundance in 16S rRNA V4 sequencing. Regions of DNA that perfectly matched *As* were, conversely, abundant throughout the experiment, although these likely reflect the presence of another species that matched the same sequence because they were consistently present in flies from all three treatments. The low abundance of the inoculated microorganisms suggests that microbes may likely to exert their selective influence during the larval stage than the adult stage, and or that their influence is through dietary modification. There is precedence that dietary modification can influence life history traits, as microbial oxidation of glucose to alpha-keto-gluconate reduces the caloric burden of the diet and the nutrients available for fat storage (49). As low-glucose diets can phenocopy the low-fat content effect of *A. tropicalis* on the flies, experiments with diets with different macronutrient ratios could be used in future experiments to test if they are the cause of compensatory adaptation in the flies. Another possible effect of the microorganisms could be heat production. Anecdotally, we observed that the lids of *As*-inoculated diets uniquely accumulated condensation before being placed in outdoor enclosures, and that poikilothermic fly populations were more active and had lower mortality in *As*-inoculated cages. If these observations are a consequence of bacterial heat production, it could suggest that the temperature of the microbe-inoculated diets might contribute to either or both of fly fitness or adaptation in a wild setting.

The reasons why development rate was subject to more consistent compensatory selection than other life history traits remain unclear. Life history theory predicts trade-offs between traits that promote reproduction (e.g., development time, fecundity) and those associated with somatic maintenance (e.g., stress resistance, lifespan). Most natural *Drosophila* populations exhibit such trade-offs, and colonization with different microorganisms can recapitulate these patterns. However, prior experimental evolution studies in *Drosophila* have shown that adaptive changes in one trait do not necessarily lead to correlated changes in others. This suggests that life history trade-offs may arise from parallel adaptation to environmental pressures rather than from direct physiological constraints. In our study, we hypothesize that host fecundity and starvation resistance were not strongly selected due to experimental conditions: flies had continuous access to diet, and egg-laying occurred over the flies’ entire reproductive period. In contrast, development time was under strong selection pressure, as flies were released into mesocosms in timed waves. Dietary effects may have further contributed to this pattern (Huang, 2015). Additionally, the observation that adult flies rarely carried the inoculated microorganisms supports the idea that microbial effects may be more pronounced during the larval stage, which lives in the diet. Collectively, these factors help explain why larval development time was more strongly selected than adult traits and suggest that disentangling microbial selection on larval versus adult phenotypes may be a promising area for future investigation

We found that changes in fly microbiota were driven in part by a notable decline in *Wolbachia* relative abundance near the conclusion of the experiment. Two previous analyses showed that *Wolbachia* prevalence tended to reduce during summer and peak in the fall across a summer-to-fall season (25) or did not change over season (45). This incongruity could partially be explained by experimental differences in each of the studies. The distinction between abundance and presence/absence could ameliorate the incongruity between our work and the work of Henry et al. (25). Lemoine et al. measured *Wolbachia* relative abundance as did we, but collected flies from true natural settings in the wild (45). By contrast, our experimental mesocosms included flies fed on natural diets with laboratory microorganisms where fly migration was blocked. The apparent variability in *Wolbachia* outcomes across studies highlights the experimental specificity of microbiota-host interactions. Indeed, prior research has reported conflicting associations (positive, negative, or neutral) between *Acetobacter* and *Wolbachia* abundance (50-53).

Taken together, our results support a model in which *D. melanogaster* adapts to its microbial environment through both ecological and evolutionary mechanisms. Microbial identity influences host fitness in the field, and host genetic adaptation can compensate for microbial effects in trait-specific ways. This study also raises several issues that identify further areas for exploration, including whether the microbiota influences adaptation dependent upon the life stage of the host (larval or adult), and whether environmental or dietary conditions contribute to or are independent of the observed compensatory adaptation. These findings underscore the importance of considering microbial colonization in studies of rapid adaptation and suggest that host-microbe interactions are a key axis of evolutionary change in natural populations.

## METHODS

### General Drosophila husbandry

General rearing conditions for *D. melanogastger* were at 25° C on a 12-h light: 12-h dark cycle at ambient atmosphere (approx. 25% humidity). Long-term *Drosophila melanogaster* stocks were maintained at 20°C on a 12-h light: 12-h dark cycle at ambient atmosphere (approx. 25% humidity) on Nutri-Fly® Bloomington Formulation Pre-Mixed Fly Food (Genesee Scientific 66-112). Bacterial strains were cultured on MRS medium (Criterion, C5932) at 30°C. *Acetobacter* strains were cultured ambient oxygen including with shaking in liquid culture and Lactic Acid Bacteria were cultured in static liquid culture to reduce oxygen exposure.

### Outdoor mescososms

Fifteen outdoor field mesocosms were placed in 2 adjacent rows of 8 and 7 mesocosms respectively on sandy soil at the University of New Hampshire Woodman Farm, Durham, NH, USA. Each mesocosm was a 2m^3^ mesh enclosure (BioQuip PO 1406C) (24, 54) supported by aluminum frames anchored to a 4” x 6” beam frame that was built around a freshly planted (< 4 weeks old from bare root dwarf stock) leafing peach tree. The mesh enclosures were secured to the wooden frame with ¼” staples, sealed with urethane spray foam, and buried in the soil to\ prevent invasion into or escape of flies from the enclosures.

A founder population of ∼15,000 flies was prepared in common garden from isofemale lines collected in Bowdoin, Maine (20 isofemale lines), Lee, NH (25 isofemale lines), and Harvard, MA (10 isofemale lines). We previously reported on the collection of these lines, which were previously confirmed to be *D. melanogaster* by morphological analysis of males and by RFLP analysis of the cytochrome C gene (55). The lines were reared for 2 generations on a molasses diet (molasses diet of 7.05% molasses (Genesee Scientific 62-117), 5.48% cornmeal (Genesee Scientific 62-100), 1.31% Brewer’s yeast, 0.74% soy flour (Genesee Scientific 62-115), 0.43% agar, 0.18% Tegosept (Genesee Scientific 20-258), and 2.91% 190 proof ethanol). Then, from each isofemale line 10 F_2_ females (relative to rearing on molasses diet) at least 2 days post eclosion were lightly anesthetized and mixed in common garden in a 1 ft^3^ *Drosophila* cage and several petri dishes of molasses diet all at 25°C and ambient humidity (∼ 25% RH), including in subsequent steps. After 2-3 days, 30 eggs were scooped from the agar surface to individual 35 ml fly vials containing 10 ml molasses diet. This population was expanded to 10,000 flies at 30 flies per vial by mixing all eclosed flies at 2-4 days post eclosion in a 1 ft^3^ cage with molasses diet, then transferring ∼30 eggs at a time to individual molasses diet vials (10 ml diet per 35 ml vial). This mixing was performed for 2 generations, and the F_4_ of the originally mixed flies reached a final population size of 15,000 adult flies. When these flies were at 4-10 days post eclosion, the flies were lightly anesthetized on CO_2_ in batches of ∼ 100 flies, and sorted into pools of 1000 flies (500 male, 500 female). Finally, each pool of 1000 flies was released into outdoor mesocosms on 28 Jun 2023.

Released flies were allowed to lay eggs on and feed *ad libitum* from aluminum loaf pans containing 400 ml of applesauce diet (1 #10 can Regal applesauce, 65.067 g Brewer’s Yeast (MP Biomedicals 903312), 1.28 L distilled water, 21.61 g agar (Genesee Scientific 66-103), 6.37 g Apex Tegosept (Genesee Scientific 20-258), 127.37 ml ethanol) administered to the mesocosms thrice weekly for the duration of the experiment. Each instance when a new loaf pan was added to the cages, the previous pan was covered with a screen mesh lid; once flies in a covered loaf pan eclosed they were released into the mesocosm by uncovering the pan, allowing flies to leave, and replacing the diet pan cover. Flies were released from diet pans 2 -3 times per week for up to 2 weeks after the first emergent flies were released, then the spent pans were discarded. Mesocosms were administered diet that had been inoculated with one bacterial treatment of either *Ao, As*, or *Wp*. The bacterial strains were streaked for isolation on solid MRS medium, grown for 48 h in liquid MRS, washed once in phosphate-buffered saline (8 g NaCl, 0.2 g KCl, 1.44 g Na_2_HPO_4_, 2.4 g KH_2_PO_4_) (PBS) and normalized in PBS to OD_600_ = 0.1. Following normalization, 2 ml was spread on the surface of the diet and incubated at room temperature for 24-72 hours.

Flies were collected from the enclosures several times throughout the experiment. For microbiome analysis, flies were collected from the mesocosms using a hand vacuum, starved for at least 1 hour to allow most microorganisms pass through with the bulk flow of diet, and frozen at -80°C. For analysis of the living fly populations, one 6 oz fly bottle containing 100 mL applesauce diet was left uncovered in each enclosure for 48 hours, then capped and shipped overnight to Provo, UT. To reduce maternal effects of field-rearing, flies were maintained under standard laboratory conditions on yeast-glucose diet for 2 generations, and F_3_ embryos of the shipped flies were reared as axenic or gnotobiotic flies in the laboratory. Collections were performed at the times shown in **FIG 1** (red arrows).

We determined that our treatments were of high fidelity (low cross-contamination) by examining the prevalence and abundance of the inoculated strains in the cages. *Ao* and *Wp* were relatively rare and low abundance in adult flies overall, but especially in mesocosms they were not inoculated to (**FIG S3**). The relatively low abundance of the strains is consistent with our observations in a previous analysis of flies over a 6.5 week period (24). When strains were detected in a mesocosm they were not inoculated to, these strains were usually not detected in all the fly samples from that mesocosm and time point (data not shown), nor were they found in earlier or subsequent time points for that mesocosm, suggesting that the detection of these reads was more likely due to contamination of sporadic samples than systemic contamination of the mesocosm. As an exception, 16S rRNA V4 reads with full-length, perfect matches to *As*, were abundant in all mesocosms. We expect that the consistent prevalence and abundance of these reads is due to natural presence of another species with an identical 16S match to *As* because the *As* reads were highly abundant from the beginning of the experiment in all the mesocosms regardless of their treatment. Alternatively, the *As* may have been broadly contaminating and phenotypic differences resulting from *Ao-* or *Wp-*inoculations may have been due to additional inoculation from *Ao* or *Wp*. When paired with the distinct phenotypes attributed to the different microbial treatments, either interpretation leads to the conclusion that different bacterial inoculations influenced fly evolution under these conditions.

Flies subjected to seasonal selection in outdoor mesocosms for 6.5 weeks were described previously (1). After the outdoor selection, flies from each mesocosm were reared in the laboratory in common garden populations for 2-3 generations, F_3_ or F_4_ embryos were derived as axenic in the laboratory, left axenic or monoassociated with *A. tropicalis* DmCS_006 or *L. brevis* DmCS_003, and their development time to adulthood and starvation resistance were measured.

### Axenic and gnotobiotic fly rearing

Embryos of *D. melanogaster* were dechorionated in 0.6% sodium hypochlorite for 5 minutes, then transferred at a target density of 30-60 embryos to 7.5 ml sterile yeast-glucose (YG) diet (10% brewer’s yeast (MP Biomedicals 903312), 10% glucose (Sigma 102506465), 1% agar (Genesee Scientific 66-103)) in 50 ml centrifuge tubes as in our usual work (56). Axenic flies were prepared with no further modification. Gnotobiotic 6-sp flies were prepared by inoculating the sterile embryos with 50 μl of a bacterial cocktail prepared by normalizing 6 separately cultured bacterial strains to OD600 = 0.1, then mixing all strains in equivalent volumes. The bacterial strains were *Acetobacter tropicalis* DmCS_006, *Acetobacter* sp. DsW_54, *Acetobacter* sp. DmW_125, *Lactiplantibacillus plantarum* DmCS_001, *Weissella paramesenteroides* DmW_115, and *Leuconostoc suionicum* DmW_098.

### Fly phenotypes

Fly population size was measured by capturing images in eight 4 ft^2^ quadrants at the top left and top right corners of each vertical side of each mesocosm. For each image a white PVC frame was held to each corner while an image was captured with a smartphone camera. Afterwards, the number of flies in each image was manually counted, and each image was treated as a different replicate. Significant differences in fly population size over time or with bacterial treatment were determined using a Kruskal-Wallis test.

Fly development times were measured 3 times per day (9AM, 1PM, 5PM) as the number of flies in each vial that had eclosed since the previous interval. Significant differences in fly development times were determined using a Cox proportional hazards model in R (57). Overall effects of time and bacterial treatment were determined by ANOVA of the model, and post-hoc Tukey comparisons were performed in R (58).

Fly starvation resistance was measured by transferring eight lightly anesthetized female flies to 30 ml vials containing 5 ml of 1% agarose. Fly survival was monitored every 4 hours until all flies in a vial were dead. Significant variation in fly survival was monitored the same as for development time.

Fly fecundity was measured by allowing a single lightly anesthetized female to lay eggs on 7.5 ml YG diet + acid preservative (0.42% propanoic acid, 0.04% phosphoric acid) in a 50 ml centrifuge tube for 18 hours. Then, the female was removed. After 10-12 days and immediately prior to eclosion of offspring, the vial was frozen at -20°C. The fecundity of each female over an 18 hour period was determined as the number of pupae that could be counted on the side of each vial. Total counts per vial were Log_10_+1 transformed and significant differences in fly fecundity between treatments were determined using a linear mixed effects model with time and bacterial treatment as fixed effects and experiment as a random effect.

Microbiota composition of gnotobiotic flies was determined by homogenizing one pools of 3 whole body male flies per vial the flies were collected from as described previously (59). Flies were sorted into 96-well plates prefilled with 100 μl PBS and 0.5 mm glass beads. Flies were mechanically lysed using a SPEX GenoGrinder for 5 min at 1750 RPM. Serial dilutions of the homogenates were made using a 96-well pipettor before 1.5 μl of each dilution was plated on MRS medium and placed in an incubator at 30° C. Colony forming units (CFUs) were counted from dilutions containing 10-40 CFU. Tan-colored colonies were classified as AAB and white or yellow colonies as LAB. CFU counts were converted to CFU fly^-1^, and significant differences in the absolute abundance of AAB, absolute abundance of LAB, and relative abundance of LAB were determined separately using a Kruskal-Wallis test.

### Preparation of microbiome libraries

DNA was extracted from pools of 3 male flies using a Zymo Research (ZR) Quick-DNA Fecal/Soil Microbe 96 Kit (D6011), following the manufacturer’s instructions. From each sampling DNA from a pool of three male flies was extracted using a Zymo Research (ZR) Quick-DNA Fecal/Soil Microbe 96 Kit (Zymo, D6011) using manufacturer’s instructions, excluding the initial Genomic Lysis Buffer preparation with β-mercaptoethanol. Briefly, flies were mechanically lysed in a SPEX GenoGrinder for 5 min at 1750 RPM and lysates were centrifuged to recover the supernatant. The supernatant was treated, washed, centrifuged, and eluted following manufacturer instructions to extract DNA. We performed PCR following established protocols using dual-barcoded primers and Accuprime Pfx polymerase (60) to amplify the V4 region of the 16S rRNA gene. PCR parameters were 95° C for 2 min, 30 cycles of 95° C for 20 s, 55° C for 15 s, and 72° C for 5 min, and a final extension of 72° C for 10 min. We confirmed successful PCR (and lack of amplification of negative controls) by gel electrophoresis of a subset of samples. Then, to normalize the samples, we divided each 20μl reaction in half, normalized each half separately using a Just-a-Plate 96 PCR Normalization and Purification Kit (Charm Biotech), then combined samples in batches of 96 samples and concentrated these using a Zymo DNA Clean & Concentrator kit. Samples were sequenced using 500 cycle Illumina chemistry on an Illumina MiSeq at the Arizona State University genomics core. Prior to sequencing, the core performed Bluepippin size selection on the amplicons in a range of 250-450 bp, and quantified and pooled the libraries in equimolar ratios.

Microbiome analysis was performed using QIIME 2 (61) on reads that passed Illumina qualify filtering. Raw 16S rRNA sequences were denoised and filtered using the DADA2 q2-dada2 plugin (62). Aside from determining the relative changes in *Wolbachia* abundance with time and treatment, amplicon sequence variants (ASVs) assigned to *Wolbachia* were filtered out for all analyses, as were reads matching non-bacterial samples, mitochondria, and chloroplasts. We created a phylogeny with fasttree2 q2-phylogeny plugin (63) and significant differences between groups were determined by PERMANOVA (64) of beta-diversity distance metrics (65, 66).Taxonomy was assigned to the ASVs using the q2-feature-classifier (67) plugin with the SILVA 138.2 OTU reference sequence database (68) using the q2-taxa plugin and the RESCRIPt package (69). Differential abundance of ASVs was determined using ANCOM (Mandal, 2015).

## Data availability

Sequences from this study will be deposited and made publicly available in the SRA at accession XXXXX when the article is accepted.

## ACKNOWLEDGEMENTS

We thank Evan Ford at the University of New Hampshire Woodman Farm and Dr. Louis Tisa for technical support. Research reported in this publication was supported in part by the National Institute of General Medical Sciences of the National Institutes of Health under Award Number R15GM140388. The content is solely the responsibility of the authors and does not necessarily represent the official views of the National Institutes of Health.

## Supporting Figures

**Fig S1.**
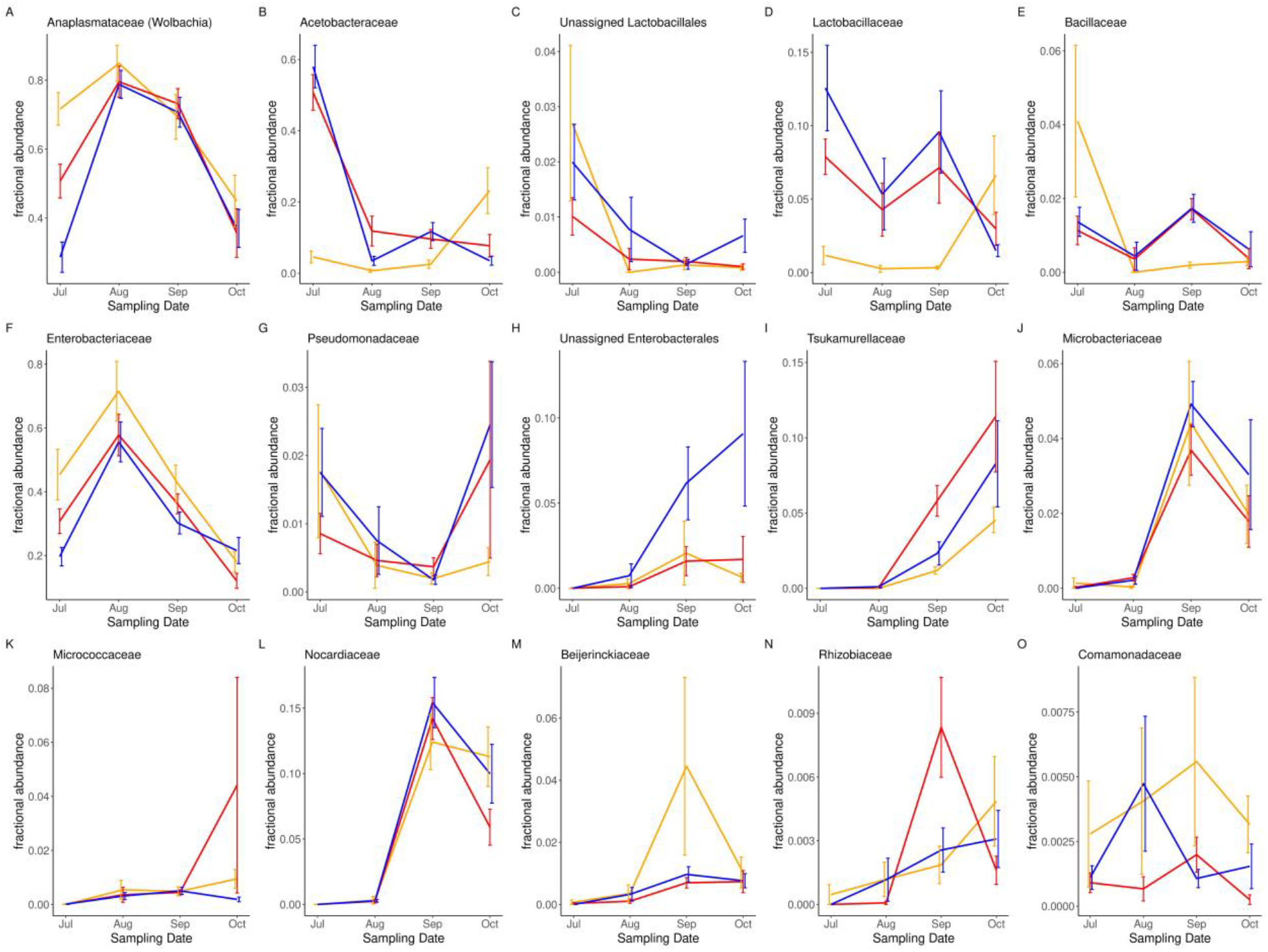
Differentially abundant ASVs. ASVs that significantly varied with time were identified using ANCOM. Analyses for *Anaplasmataceae* (*Wolbachia*) (A) was performed using an OTU table where *Wolbachia* reads were included and which was rarefied to 1000 reads per sample (**DATA S1**). All other taxa were identified from an OTU table where *Wolbachia* reads were exluded and rarefied to 537 reads per sample (**DATA S2**). ASVs in D) and H) also varied significantly with the microbial treatment. Line colors represent mesocosms inoculated with *Ao* (red), *As* (orange), or *Wp* (blue).

**Fig S2.**
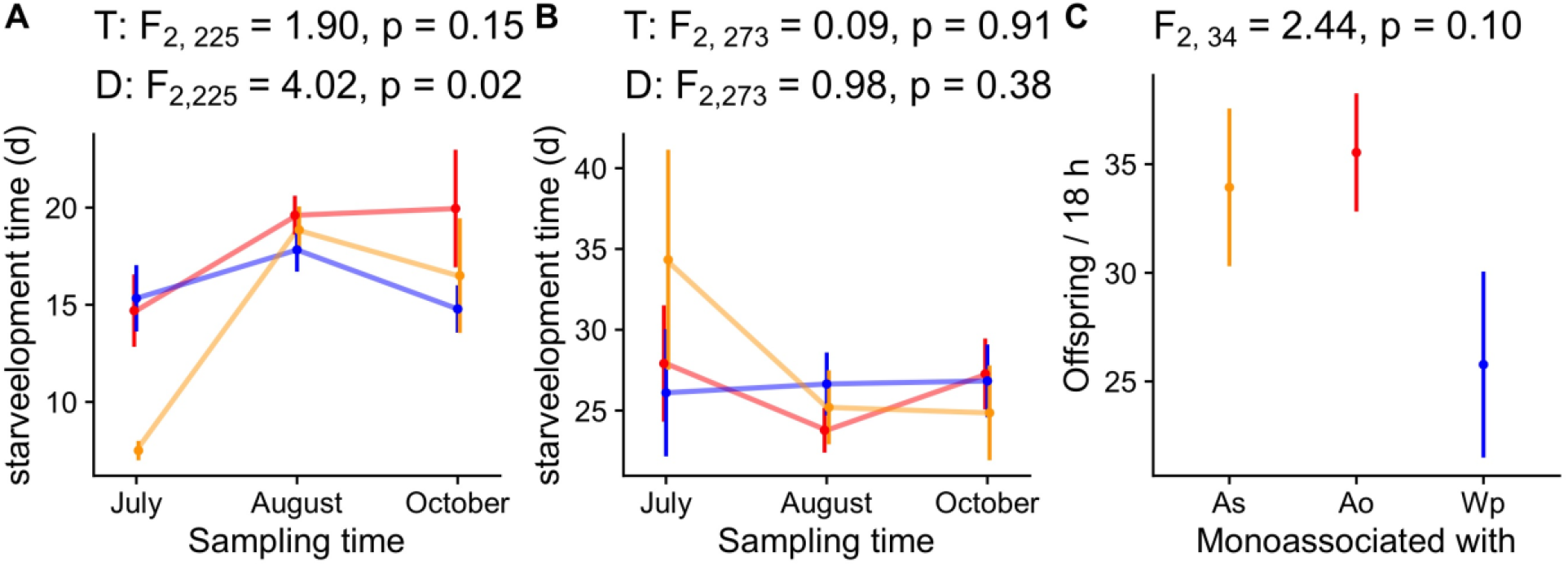
Fecundity of seasonally-selected *D. melanogaster*. The number of offspring produced by a single female in 18 hours was measured in A) axenic or B) gnotobiotic flies taken from the field and reared in the laboratory after several generations of common garden, or C) F3 offspring of the starting populations in the cages, monoassociated with the treatment strains in the laboratory.

**Fig S3.**
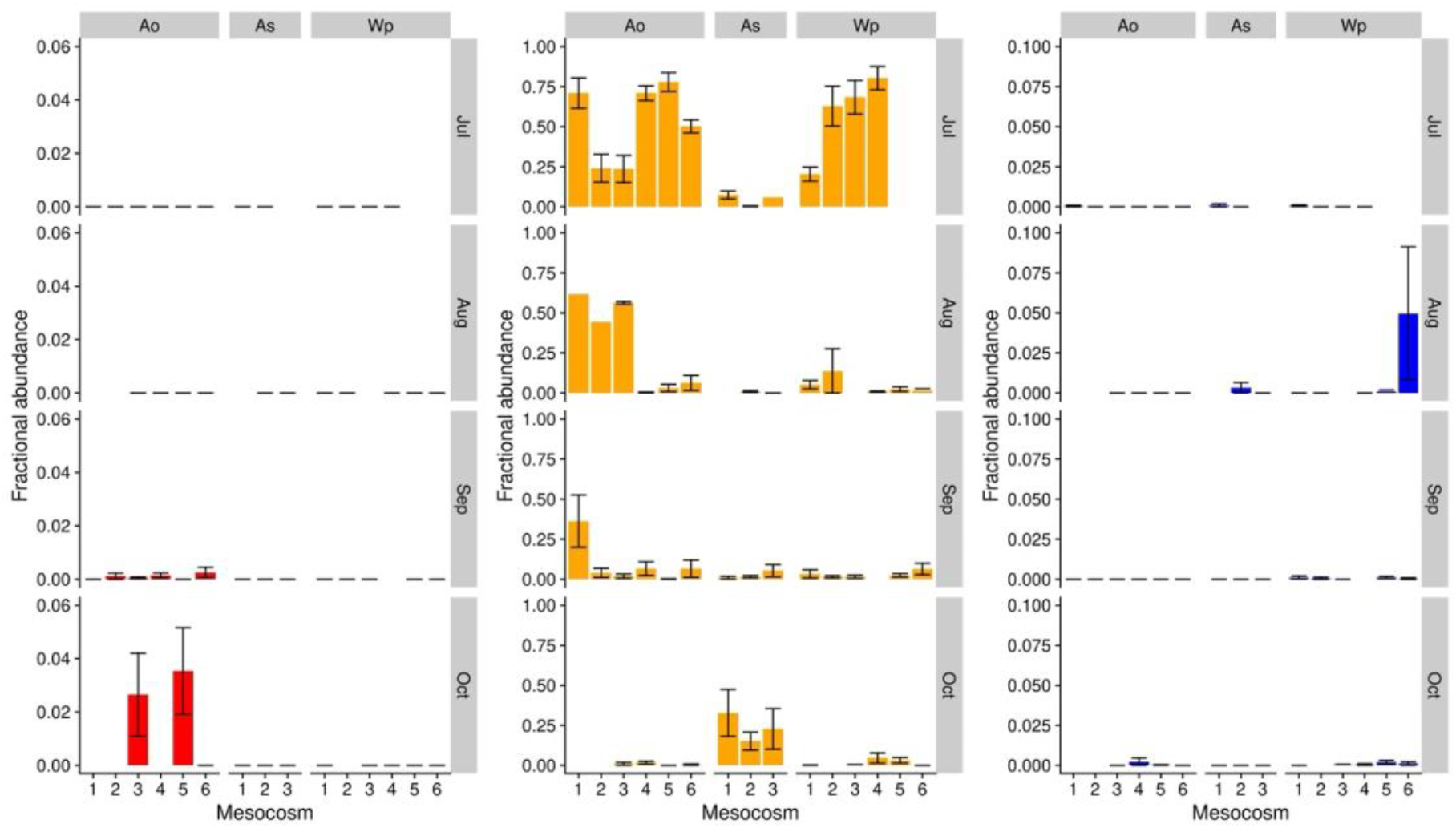
Detection of inoculated strains throughout the experiment. Flies were inoculated with *Ao, As*, or *Wp* over the course of four months (Jul-Oct). Separate panels for the fractional abundance of *Ao* (red), *As* (orange), and *Wp* (blue) show the presence of ASVs with full-length 100% matches to the inoculated treatment strains.

**Table S1.** PERMANOVA results.

|  |  | Bray-Curtis |  |  |  |  | Unweighted Unifrac |  |  |  |  | Weighted Unifrac |  |  |  |
| --- | --- | --- | --- | --- | --- | --- | --- | --- | --- | --- | --- | --- | --- | --- | --- |
|  | df | SS | R<sup>2</sup> | F | p |  | SS | R<sup>2</sup> | F | p |  | SS | R<sup>2</sup> | F | p |
| Bacteria | 2 | 1.66 | 0.02 | 3.78 | <10<sup>-3</sup> |  | 0.73 | 0.02 | 3.01 | <10<sup>-3</sup> |  | 0.44 | 0.02 | 3.52 | <10<sup>-3</sup> |
| Date | 3 | 18.95 | 0.2 | 28.69 | <10<sup>-3</sup> |  | 8.21 | 0.17 | 22.51 | <10<sup>-3</sup> |  | 6.26 | 0.23 | 33.43 | <10<sup>-3</sup> |
| Bacteria * Date | 6 | 4.23 | 0.05 | 3.21 | <10<sup>-3</sup> |  | 1.09 | 0.02 | 1.49 | 0.01 |  | 1.44 | 0.05 | 3.84 | <10<sup>-3</sup> |
| Residual | 305 | 67.13 | 0.72 |  |  |  | 37.08 | 0.78 |  |  |  | 19.02 | 0.69 |  |  |
| Total | 317 | 93.3 | 1 |  |  |  | 47.4 | 1 |  |  |  | 27.65 | 1 |  |  |

